# CRISPR/Cas9 screens implicate RARA and SPNS1 in doxorubicin cardiotoxicity

**DOI:** 10.1101/2022.08.01.502373

**Authors:** Wenjian Lv, Yeng Shao, Atsushi Hoshino, Zoltan Arany, Kiran Musunuru, Chris McDermott-Roe

## Abstract

Doxorubicin (DOX) is an efficacious chemotherapy compound used to treat various cancers which elicits severe side effects, including heart failure. Uptake of DOX by cardiomyocytes causes metabolic dysfunction and cell death but causal mechanisms remain largely undefined. We applied genome-wide CRISPR/Cas9 knockout screens to discover genetic modifiers of DOX-induced cardiomyocyte cell death, and independently, DOX uptake and clearance. Both screens discovered known and novel factors. In cell death screens and validation studies, loss of retinoic acid receptor-*α* (RARA) predisposed cardiomyocytes to DOX-mediated cell death. Conversely, RARA activation reduced DOX cytotoxicity in wild type cardiomyocytes. RNA-Seq analysis revealed that whilst DOX caused large-scale suppression of metabolic and mitochondrial gene expression, RARA activation mitigated this effect. In DOX accumulation screens, an essential role for lysosomes in DOX clearance was observed. Loss of Sphingolipid Transporter 1 (SPNS1) led to DOX hyperaccumulation, suppression of autophagy, increased DNA damage, and increased cell death. Hence, SPNS1 plays a key role in buffering against DOX accumulation and toxicity. Collectively, our study nominated hundreds of drug-gene interactions, providing a springboard for exploration of causal mechanisms, and a technical framework for future screening campaigns.

## Introduction

Anthracycline chemotherapy is a cornerstone of the anti-cancer regimen, particularly for childhood cancer and breast cancer, in which average 5-year survival rates are currently >85% (1,2). However, the clinical utility of anthracyclines such as doxorubicin (DOX) is offset by adverse effects, notably cardiotoxicity, which can range in severity from an asymptomatic decline in ejection fraction to heart failure (3). Anthracycline cardiotoxicity is common, affecting an estimated 9-18% of patients, and symptoms can occur anytime from commencement of chemotherapy to months or even years after completion (4). With increased use and improved survival rates, the population of individuals at risk of developing anthracycline cardiotoxicity secondary to chemotherapy is expanding.

A major hurdle in the field of cardio-oncology is a lack of knowledge regarding the mechanisms which cause anthracycline cardiotoxicity. Production of reactive oxygen species (ROS) is regarded as a key disease-causing mechanism (the so-called ‘ROS hypothesis’ (5)) but the therapeutic benefit of antioxidants remains unclear (6). The search for additional therapeutically-targetable mechanisms is ongoing. Recent studies have spotlighted additional important liabilities, such as heightened TOP2B expression (7), and revealed the pleiotropic nature of DOX cardiotoxicity (8). Indeed, myriad pathological processes have been implicated in DOX cardiotoxicity, including oxidative damage, mitochondrial dysfunction, apoptosis, impaired autophagy, and calcium mishandling. However, it remains unclear how these pathological processes are triggered and if disease is driven by a single event or multiple events operating simultaneously.

Reverse genetics-based approaches, including candidate gene association studies and more recently, genome-wide association studies (GWAS), have provided important insights into anthracycline cardiotoxicity. The first such investigation, by Wojnowski and colleagues (9), revealed the importance of DOX efflux transporters and ROS production, and in each case identified susceptibility polymorphisms. Recently, a GWAS of childhood cancer survivors found a common polymorphism in the Retinoic acid receptor-γ (RARG) gene (rs2229774; S427L) confers increased susceptibility to anthracycline cardiotoxicity risk (10). Forward genetics screening offers a complementary means to define causal mechanisms and circumvents *a priori* assumptions, logistical complications, and statistical rigor considerations inherent to many reverse genetics studies. The first report assayed DOX tolerance in a large panel of yeast strains and described homologous recombination, transcriptional regulation, chromatin remodeling, and cell stress as vulnerable pathways (11). An shRNA screen in lymphoma cells identified Top2a, the DOX target gene, as well as Chk2 and p53 as important determinants of DOX toxicity (12). Linkage analysis in lymphoblastoid cell lines treated with daunorubicin revealed that multiple naturally occurring polymorphisms, including those in INPP4B and CDH13, modify drug toxicity (13).

In this study, we adopted a forward genetic approach, applying genome-wide and targeted CRISPR/Cas9 screens across various cardiomyocyte-derived cell lines to probe for genes, proteins, and pathways involved in DOX cardiotoxicity. For phenotypic readouts, we analyzed cell death and DOX accumulation, both of which drive cardiotoxicity *in vitro* and *in vivo* (14,15). Collectively, we discovered hundreds of drug-gene interactions, underscoring the crucial interplay between genetic background and drug sensitivity. We went on to confirm the role of Retinoic acid receptor-*α* (RARA) status as a determinant of DOX sensitivity and Sphingolipid Transporter 1 (SPNS1) in facilitating DOX clearance.

## Results

### Genome-wide CRISPR/Cas9 screening of modifiers of doxorubicin-mediated cell death

We first sought to define genetic modifiers of doxorubicin (DOX)-mediated cardiomyocyte death, a key contributor to DOX-mediated cardiotoxicity (16), using genome-wide CRISPR/Cas9 screens. An immortalized mouse cardiomyocyte cell line, HL-1 (17), was transduced with GeCKOv2 lentiCRISPR library (18), subjected to antibiotic selection, expanded, and treated with either DOX to kill ~90% of the culture (1μM for 3 days) or DMSO. The remaining cells were recovered and genomic DNA was deep-sequenced to measure sgRNA representation (Figure 1A). In theory, sgRNAs enriched in DOX-treated cells versus DMSO-treated cells specify genes whose perturbation desensitize cells to DOX (i.e., reduce cell death). Conversely, depleted sgRNAs specify genes whose perturbation sensitize cells to DOX (i.e., increase cell death). Using MAGeCK (19), we discovered 213 enriched sgRNAs targeting 36 genes and 506 depleted sgRNAs targeting 87 genes (false discovery rate (FDR)<0.25) (Figure 1B). Validating our approach, loss of Top2A, the primary DOX target gene (20), represented the strongest desensitizing hit (log_2_ fold change (log_2_FC)=2.1, *P*=2.2×10^-7^, FDR<0.001) whilst loss of Abcb1b, a DOX efflux transporter (21), was the strongest sensitizing hit (log_2_FC=-1.9, *P*=2.2×10^-7^, FDR<0.001). Gene ontology (GO) term analysis revealed over-representation of pathways relevant to DOX cardiotoxicity, including the DNA damage response and programmed cell death (Figure 1C). To validate the screen, we disrupted a panel of genes and used an orthogonal assay to measure DOX toxicity (Supplementary Figure 1). Relative to wild type HL-1 cells and HL-1 cells transduced with a non-targeting sgRNA, most KO cell lines exhibited altered sensitivity to DOX consistent with our screen. A second exploratory screen was performed in a human cardiomyocyte cell line, AC16 (22), using the same strategy (Figure 1A). We discovered 251 enriched sgRNAs targeting 69 desensitizing genes and 100 depleted sgRNAs targeting 25 sensitizing genes (Figure 1D). As above, the desensitizing and sensitizing populations contained canonical regulators of DOX-mediated cell death, and GO term analysis revealed over-representation of relevant pathways, including intrinsic apoptosis, P53 signaling, and non-homologous endjoining (Figure 1E).

**Figure 1:**
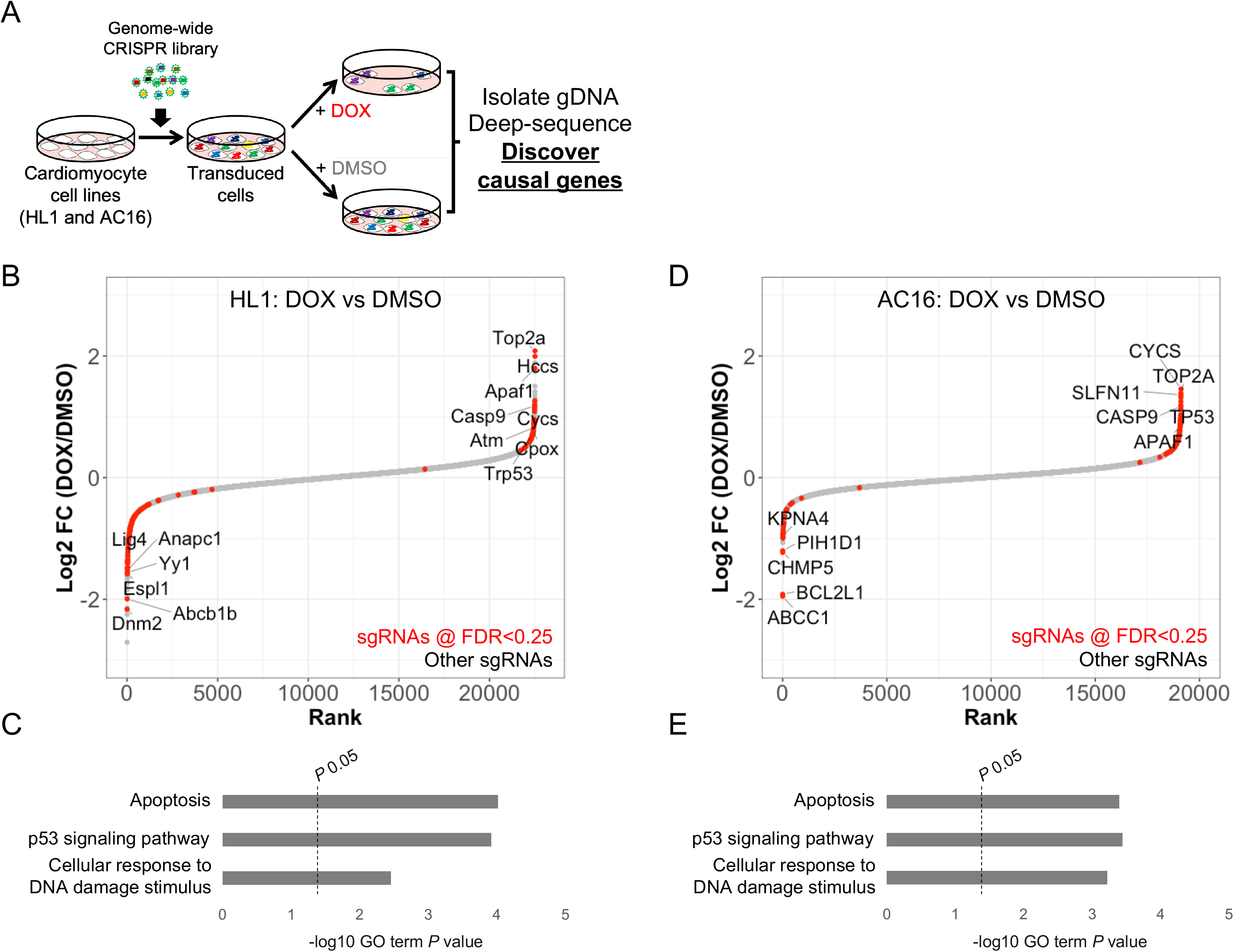
Defining genetic modifiers of doxorubicin cytotoxicity via exploratory genome-wide CRISPR/Cas9 knockout screens in HL-1 mouse cardiomyocyte and AC16 human cardiomyocyte cell lines. (A) Experimental approach. HL-1 and AC16 cells were transduced with genome-wide CRISPR/Cas9 knockout libraries at ~300x coverage, expanded, and treated with either doxorubicin (DOX) (1μM for 3 days) or DMSO, after which sgRNA representation was determined via next-generation sequencing. (B and D) Ranked dot plots showing relative abundance of sgRNAs at gene-level in HL-1 (B) and AC16 (D) cells treated with DOX versus DMSO. Each point represents a gene. Red points are those differentially-represented in DOX-treated cells versus DMSO-treated cells (false discovery rate (FDR)<0.25). Select hits are labelled. (C and E) Top gene ontology (GO) terms and *P* values obtained via DAVID (see Methods) associated with perturbations leading to DOX desensitization in HL-1 (C) and AC16 (E) cells.

Collectively, our HL-1 and AC16 CRISPR screens identified 217 high confidence gene-drug interactions (FDR<0.25) and provided insights into the molecular pathobiology of DOX cardiotoxicity. In both screens, the highest scoring genes are established regulators of intrinsic apoptosis, which tallies with recent work (23), underscoring the regulated nature of DOX-mediated cell death. Of note, most non-apoptosis hit genes possessed moderate effect sizes and spanned a wide functional spectrum. This suggests causal mechanisms are numerous, functionally diverse, triggered in a concurrent fashion, and converge on apoptosis to induce cell death.

We next aimed to validate our findings in human iPSC-derived cardiomyocytes (iCMs), rationalizing that a targeted CRISPR KO screen assaying a prioritized subset of genes with an expanded panel of sgRNAs at higher coverage would strengthen our capacity to identify robust, physiologically-relevant modifier genes. From each exploratory screen, we selected the top 500 modifier genes and created a targeted CRISPR-Cas9 KO library comprising ~7800 unique sgRNAs targeting 952 unique genes (8 sgRNAs per gene) and >200 non-targeting sgRNAs. The library was transduced at high coverage (x1000) into a healthy iPSC line (SV20,24), which was then differentiated into iCMs via the standard Wnt modulation protocol with lactate selection (25,26). Mutagenized iCMs were treated with a half-maximal lethal dose of DOX (LD_50_; 250nM for 5 days) or DMSO, after which live cells were recovered, genomic DNA isolated, and sgRNA representation analyzed as described (and see Methods) (Figure 2A). The experiment was conducted in biological duplicate using independently differentiated batches of iCMs to maximize robustness. Of the 7601 gene-specific sgRNAs in our targeted library, 1047 (~14%) were differentially represented by 2-fold or more in DOX-treated iCMs versus DMSO-treated iCMs compared to 12 of the 200 non-targeting sgRNAs (6%) (Fisher’s exact two-tailed *P*=0.0008, Supplementary Figure 2). Using a modified inclusion threshold to legislate for variable sgRNA performance (see Methods), we nominated 11 and 35 high confidence DOX-sensitizing and DOX-desensitizing perturbations, respectively. The most significantly depleted sgRNAs were those targeting the retinoic acid receptor-*α* (RARA) gene indicating that RARA loss-of-function exacerbates DOX-mediated cell death (Figure 2B). To confirm this, DOX toxicity was examined in an independent iPSC line (hB53,27) in which we first disrupted RARA via CRISPR (Figure 2C). Consistent with our screen, RARA KO iCMs were sensitized to DOX compared with isogenic control iCMs (Figure 2D). Given that RARA deficiency exacerbated DOX cell death, we explored whether its activation imparts protection. Tamibarotene (TBT) is an RARA agonist used for treatment of acute myeloid leukemia in cases of all-trans retinoic acid (ATRA) resistance (28). In iCMs pre-/co-treated with TBT, DOX-mediated cell death was significantly lower compared with cells treated with DOX alone (Figure 2E). Additionally, in iCMs treated with DOX (1μM for 24-hours) plus TBT, myofibrillar disarray was less severe compared with iCMs treated with DOX alone (Figure 2F). RARA is an important transcription factor so we tested if protection stemmed from changes in gene expression. RNA-Seq was conducted on iCMs treated with (i) DMSO, (ii) DOX, (iii) TBT plus DOX, and (iv) TBT (4 biological replicates per condition). Significant transcriptional changes were observed across all conditions as evident from principal components analysis and pairwise comparisons of transcript abundance (Figure 3A,B). Notably, in the DOX vs DMSO comparison, almost all detected mRNAs (11087 of 12654) were differentially-expressed (FDR<0.05) and unbiased gene set enrichment analysis (GSEA) revealed the down-regulated population was highly enriched with genes encoding mitochondrially localized proteins. Conversely, in the TBT plus DOX vs DOX comparison, whilst transcriptional changes were subtle, mitochondrial genes were again highly enriched, but this time in the up-regulated population (Figure 1C,D, Supplementary Figure 4). Hence, we suggest TBT imparts protection by buffering against DOX-mediated suppression of mitochondrial gene expression, and in doing so alleviating metabolic dysfunction.

**Figure 2:**
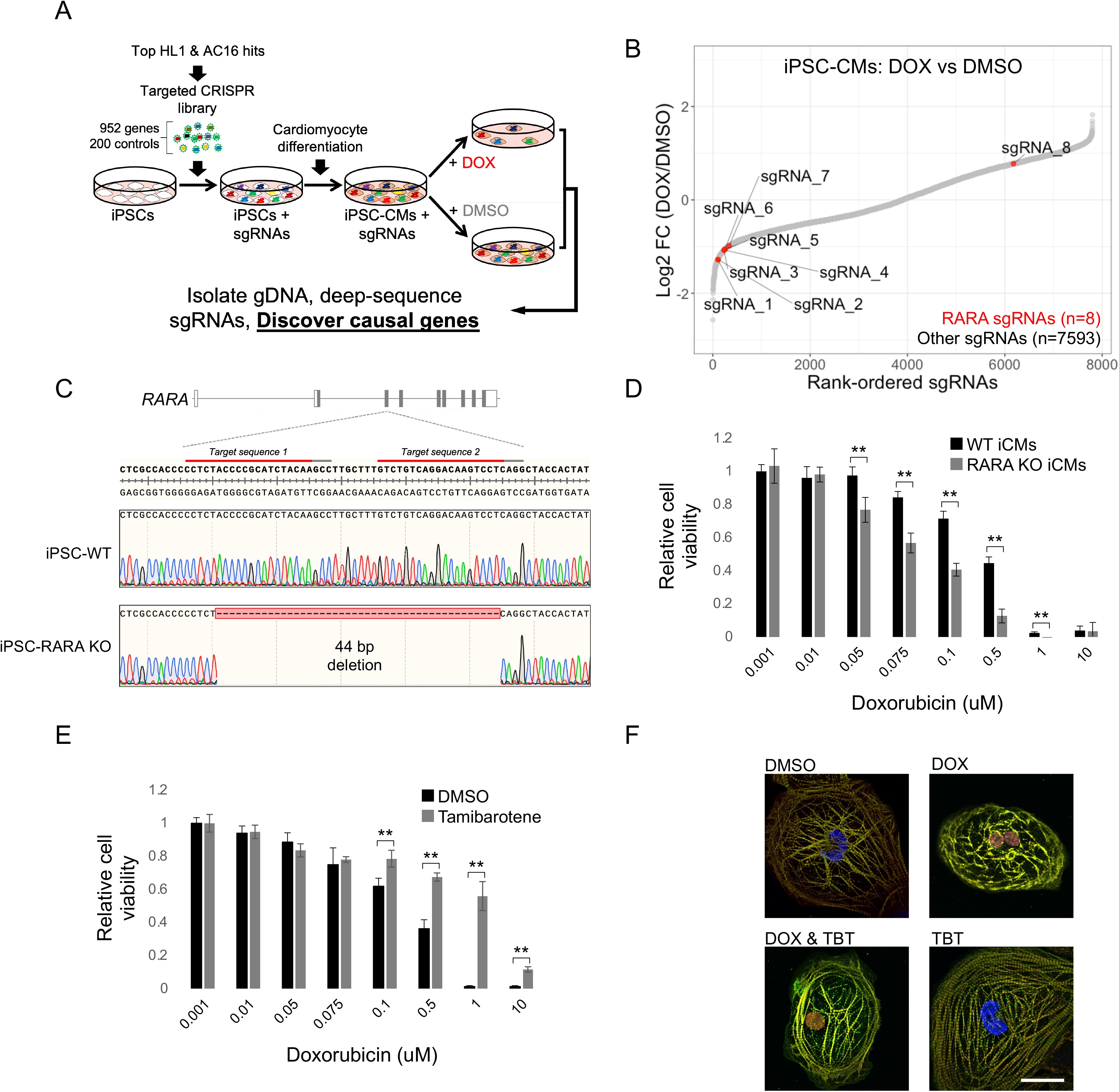
Validation of exploratory screening data using targeted CRISPR/Cas9 knockout screens in human iPSC-derived cardiomyocytes and discovery of RARA status as a determinant of doxorubicin cardiotoxicity. (A) Experimental approach. The top hits from the HL-1 and AC16 screens were used to design a targeted CRISPR/Cas9 knockout (KO) library, which was transduced into iPSCs at >1000x coverage. Following differentiation, iCMs were treated with doxorubicin (DOX) or DMSO. sgRNA representation was determined via next-generation sequencing. (B) Ranked dot plot showing relative abundance of sgRNAs in iCMs treated with DOX versus DMSO. Each point represents an sgRNA. Red points are sgRNAs targeting the RARA gene. (C) CRISPR-based RARA KO in a healthy iPSC line. Two sites in exon 3 were targeted simultaneously. Multiple clones bearing single and double allelic indels were recovered, including the homozygous 44 bp deletion mutant shown. (D) Wildtype (WT) and RARA KO iPSCs were differentiated into iCMs and treated with specified concentrations of DOX for 72-hours. Cell viability was measured via alamarBlue assay (see Methods). (E) Effect of Tamibarotene (TBT) pre-/co-treatment on DOX toxicity in wildtype iCMs. Wildtype iCMs were pre-treated for with 1μM Tamibarotene or DMSO for 24 hours and then treated with DOX in the presence of Tamibarotene (1μM) or DMSO. (F) Myofibrillar dis/organization in WT iCMs treated with DOX, DOX plus TBT, TBT, and DMSO. Red, cardiac troponin T; green, α-actinin; blue, DAPI. Scale bar: 20μm.

**Figure 3:**
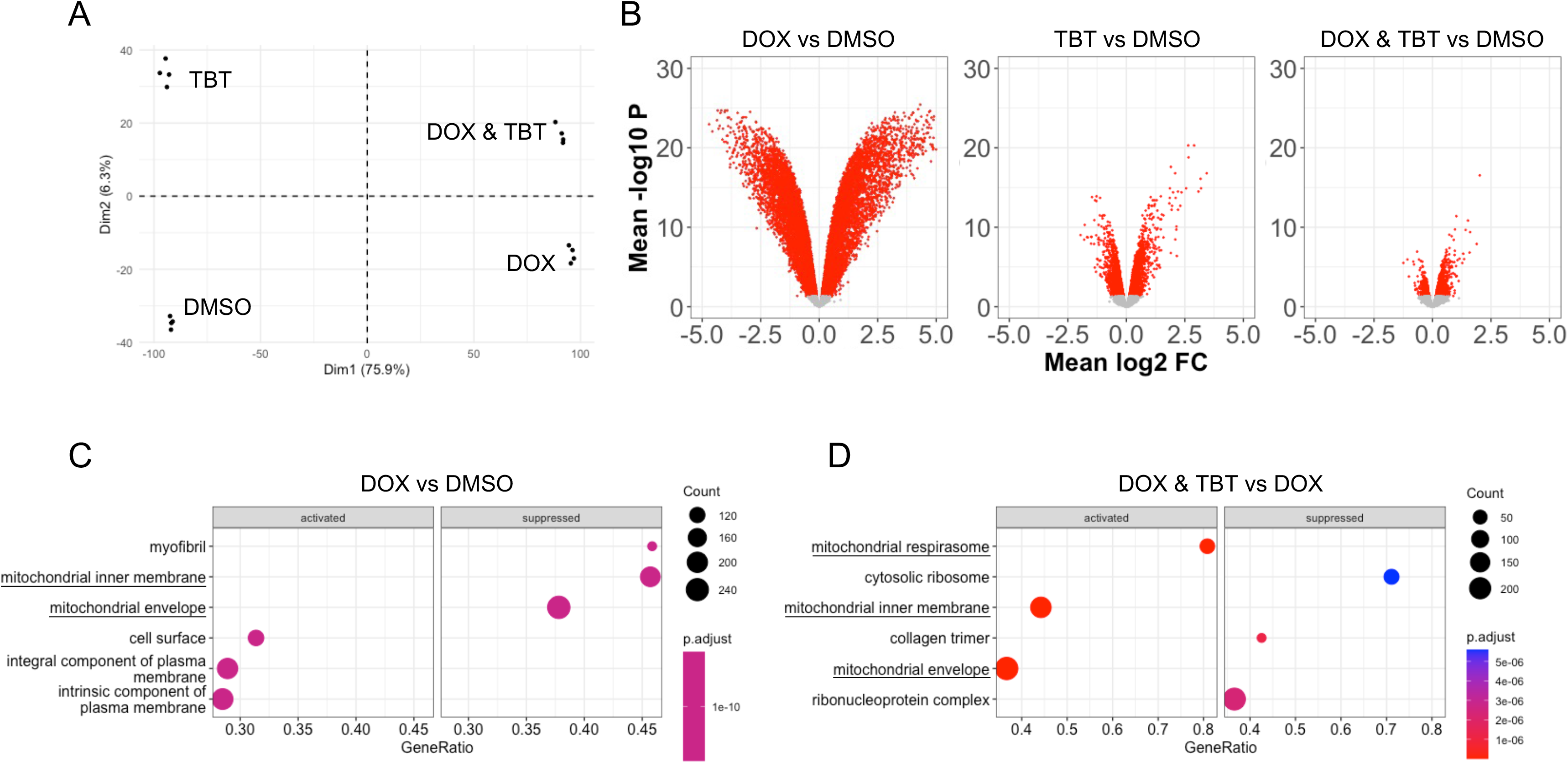
Tamibarotene mitigates doxorubicin-mediated suppression of mitochondrial gene expression. (A) Principal components analysis (PCA) plot based on transcript profiles in iCMs treated with doxorubicin (DOX), Tamibarotene (TBT), DOX plus TBT, and DMSO. (B) Pair-wise volcano plots of RNA-Seq data. Each point represents a transcript. Red points denote those differentially-expressed (false discovery rate (FDR)<0.05). (C) Gene ontology (GO) term analysis showing top cellular component terms associated with up- and down-regulated transcripts (activated and suppressed, respectively) in iCMs treated with DOX compared with iCMs treated with DMSO. (D) As in C, but in iCMs treated with DOX & TBT compared with iCMs treated with DOX.

### Discovery of genetic modifiers of doxorubicin (DOX) accumulation

Persistence of DOX within cardiomyocytes is another crucial toxicity mechanism (29–32). We next applied CRISPR screening for discovery of factors involved in its uptake and clearance. To test feasibility, we incubated AC16 cells with DOX for 24-hours and observed a robust and dose-dependent increase in fluorescence intensity by flow cytometry (Figure 4A). Additionally, AC16 cells lacking ABCC1, a known DOX efflux pump (9), exhibited increased fluorescence relative to wild type AC16 cells following DOX incubation (Figure 4B). Hence, flow cytometry enables measurement of DOX accumulation and is sensitive enough to detect an increase resulting from loss of a known DOX efflux factor. We twinned CRISPR/Cas9 mutagenesis with fluorescence-activated cell sorting (FACS) to screen for determinants of DOX accumulation genome-wide. AC16 cells were transduced with the Brunello genome-wide CRISPR/Cas9 library and incubated with 1μM DOX for 24-hours, which led to DOX uptake without overtly impacting cell health. Cells were then sorted into two bins via FACS; those with low levels of DOX (the bottom 15%) and those with high levels of DOX (the top 15%) (Figure 4C and Supplementary Figure 3). Cells with perturbations reducing DOX uptake (or increasing clearance) theoretically fall in the low DOX bin whilst cells with perturbations increasing uptake (or reducing clearance) fall in the high DOX bin. We identified 125 enriched sgRNAs targeting 32 unique genes in the low DOX bin and 84 enriched sgRNAs targeting 23 unique genes in the high DOX bin (FDR<0.25) (Figure 4D,E). GSEA of perturbed genes in the low DOX bin identified ribosome biogenesis and translation as top-scoring functions. Meanwhile, the high DOX bin was enriched for perturbations in genes involved in lysosome function and V-type ATPase complex-mediated proton transport (Supplementary Figure 5). Collectively, the screen identified over 50 high confidence drug-gene interactions and implicated various molecular functions, indicating that like DOX-mediated cell death, uptake and clearance of DOX are regulated by fundamental cellular processes.

**Figure 4:**
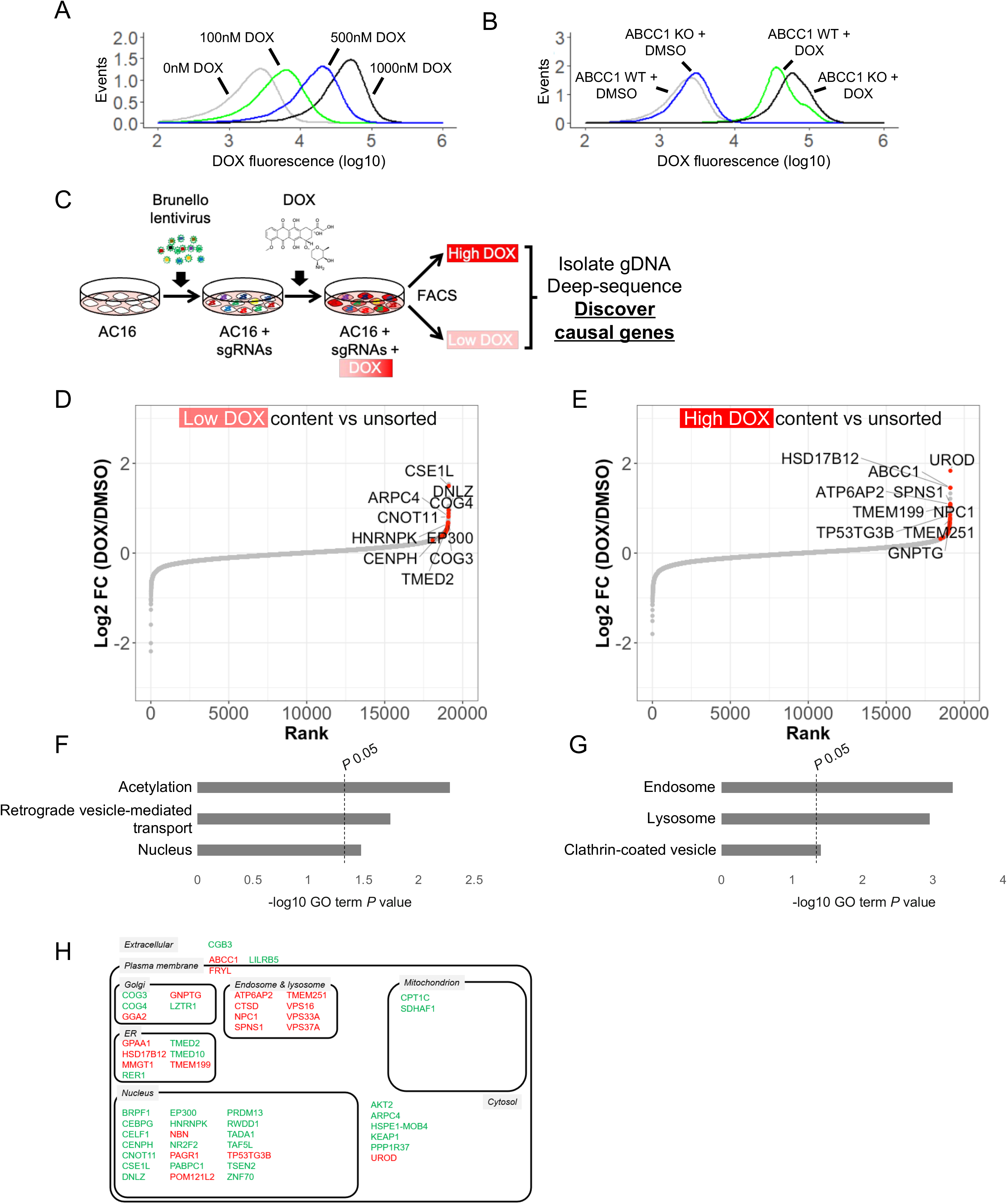
Discovery of factors regulating doxorubicin accumulation via genome-wide CRISPR/Cas9 knockout screens in AC16 human cardiomyocytes. (A) Incubating AC16 cells with increasing amounts of doxorubicin (DOX) correlates with increased fluorescence (585nm) measured by flow cytometry. (B) ABCC1 knockout (KO) AC16 cells accumulate more DOX than ABCC1 wildtype (WT) cells. (C) Experimental approach. AC16 cells were transduced with a genome-wide CRISPR/Cas9 knockout (KO) library at ~300x coverage, expanded, and treated with DOX (1μM for 1 day), after which cells were separated by FACS based on DOX accumulation into a low content bin (bottom 15%) and a high content bin (top 15%). sgRNA representation was measured via next-generation sequencing. (D and E) Ranked dot plots showing relative abundance of sgRNAs at gene-level in low content bin (D) and high content bin (E) versus unsorted cells. Each point represents a gene. Red points are those differentially-represented (false discovery rate (FDR)<0.25). Select hits are labelled. (F and G) Top gene ontology (GO) terms and *P* values obtained via DAVID (see Methods) associated with perturbations leading to low (F) and high (G) DOX accumulation in AC16 cells. (H) Cellular localization of all hit genes at FDR<0.25.

Genes in the high DOX bin likely function in the catabolism and/or efflux of DOX since their perturbation increased accumulation. The joint-top hit was ABCC1 (log_2_ fold change (log_2_FC)=1.5, *P*<3×10^-7^, FDR<0.001), a known DOX efflux pump and cardiotoxicity susceptibility gene (9), examined in our feasibility studies. Most other top hits in the high DOX bin reside within or regulate endosome/lysosome function (Figure 1G). Strikingly, every lysosome gene which scored as a hit was found in the high DOX bin. Equally interesting was the lack of canonical autophagy regulators given the documented, albeit complex, link between autophagy and DOX cardiotoxicity (33–38). Indeed, over-expressing a Beclin 1 mutant which hyperactivates autophagy (39) did not significantly affect DOX accumulation in AC16 cells (Supplementary Figure 6).

Sphingolipid Transporter 1 (SPNS1), joint-top hit in the high DOX bin (log_2_FC=1.1, *P*<3×10^-7^, FDR<0.001), encodes a lysosomal protein believed to regulate carbohydrate transport and autophagic lysosome reformation (40). SPNS1 was disrupted in AC16 cells using an independent sgRNA (Figure 5A). Validating our screen, AC16-SPNS1 KO cells accumulated substantially more DOX than AC16 cells transduced with a control (con) sgRNA (Figure 5B). Real-time confocal microscopy of DOX-treated cells enabled visualization of DOX accumulation (Figure 5C and Supplementary Figure 7). In AC16-con cells, DOX was detected in the nucleus, throughout the cytosol, and to a lesser extent in perinuclear vesicles. In AC16-SPNS1 KO cells, a significant fraction of DOX was present in perinuclear bodies, which we hypothesized were lysosomes. To test this, cells were treated with DOX followed by LysoTracker Green (LTG) and viewed by fluorescence microscopy. In AC16-con cells, discernible LTG positive bodies were scarce and there was limited overlap between DOX and LTG. In contrast, AC16-SPNS1 KO cells exhibited greater LTG signal and co-localization with DOX was readily apparent (Figure 5D). SPNS1 deficiency also affected distribution of LC3, a commonly used marker of autophagic flux (Figure 5E). In AC16-SPNS1 KO cells, we observed a marked increase in LC3 puncta following DOX treatment compared with AC16-con cells, although LC3 punta did not robustly co-localize with DOX (data not shown). The role of SPNS1 in cardiomyocytes has not been examined so we disrupted the gene in a healthy donor-derived iPSC line (27) via CRISPR (Figure 6A) and differentiated into iCMs as described above. Mirroring our AC16 studies, DOX accumulation was elevated in iCM-SPNS1 KO cells compared with iCM-con cells as assessed by flow cytometry and fluorescence microscopy (Figure 6B,C). DOX accumulation was further exacerbated by Torin 1 in iCM-SPNS1 KO cells but not in iCM-con cells (Figure 6F) suggesting loss of autophagic homeostatic control contributes to DOX accumulation. This was further examined by measuring lysosome abundance at baseline and following nutrient starvation (Figure 6G). In iCM-con cells, lysosome content was unchanged in starved cells compared with fed cells but was markedly increased in iCM-SPNS1 KO cells, suggesting that, akin to kidney epithelial cells (40), SPNS1 loss-of-function prevents autophagic lysosome reformation in human cardiomyocytes. This is a likely cause of the increase in DOX accumulation. We next investigated DOX accumulation in more detail. DOX was detected in nuclei and perinuclear punctate bodies in iCM-con cells (Figure 6C). The latter was confirmed to be lysosomes based on LAMP1 co-localization (Supplementary Figure 8). Using image cytometry, we observed punctate, perinuclear DOX deposits which were larger and more numerous in iCM-SPNS1 KO cells than in iCM-con cells (Figure 6D). DOX was apparent for weeks following its removal in iCM-con and iCM-SPNS1 KO cells (data not shown), indicating a limited capacity to clear DOX. Consistent with DOX blocking autophagy, lipidated LC3 (LC3-II) abundance increased in AC16-con cells following DOX treatment and more so in iCM-SPNS1 KO cells (Figure 6E). As reflected by phosphorylation of H2AX, DOX-mediated DNA damage was also enhanced in iCM-SPNS1 KO cells compared with iCM-con cells (Figure 6E). Finally, we investigated if SPNS1 deficiency affects DOX toxicity. Following DOX exposure, cell viability in iCM-SPNS1 KO cells was significantly reduced compared with iCM-con cells (Figure 6G). Collectively, these data show that SPNS1 plays an important role in cardiomyocytes by buffering against DOX accumulation, presumably via lysosome recycling, limiting autophagic dysfunction and DNA damage, and ultimately mitigating DOX toxicity.

**Figure 5:**
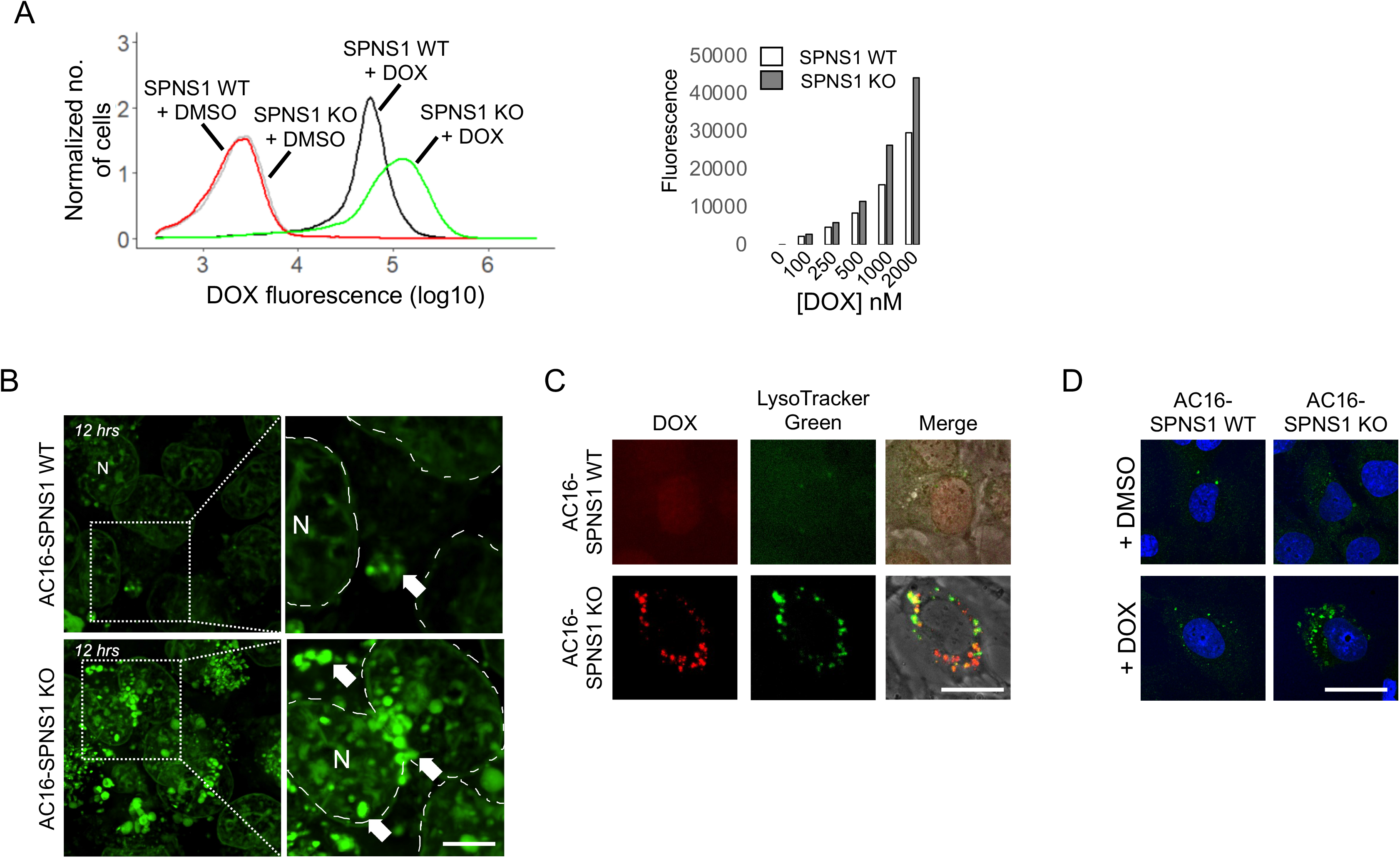
Validation of SPNS1 as a modifier of DOX accumulation. (A) Flow cytometry analysis of doxorubicin (DOX) accumulation in SPNS1 wildtype (WT) and SPNS1 knockout (KO) AC16 cells. Representative flow cytometry histogram (left) and background-corrected median fluorescence intensities following incubation with specified concentrations of DOX (right). (B) Time-lapse confocal microscopy of DOX accumulation in SPNS1 WT and SPNS1 KO AC16 cells. 12-hour timepoint shown. Dashed lines demarcate nuclear membranes for clarity; N, nucleus; arrows specify perinuclear DOX aggregates. Scale bar: 5μm. (C) Live cell fluorescence imaging of SPNS1 WT and SPNS1 KO AC16 cells following DOX treatment (1μM for 24-hours) and LysoTracker Green (75nM for 1-hour). Scale bar: 20μm. (D) Fluorescence imaging of LC3 distribution in SPNS1 WT and SPNS1 KO AC16 cells following DOX treatment (1μM for 24-hours). Scale bar: 20μm.

**Figure 6:**
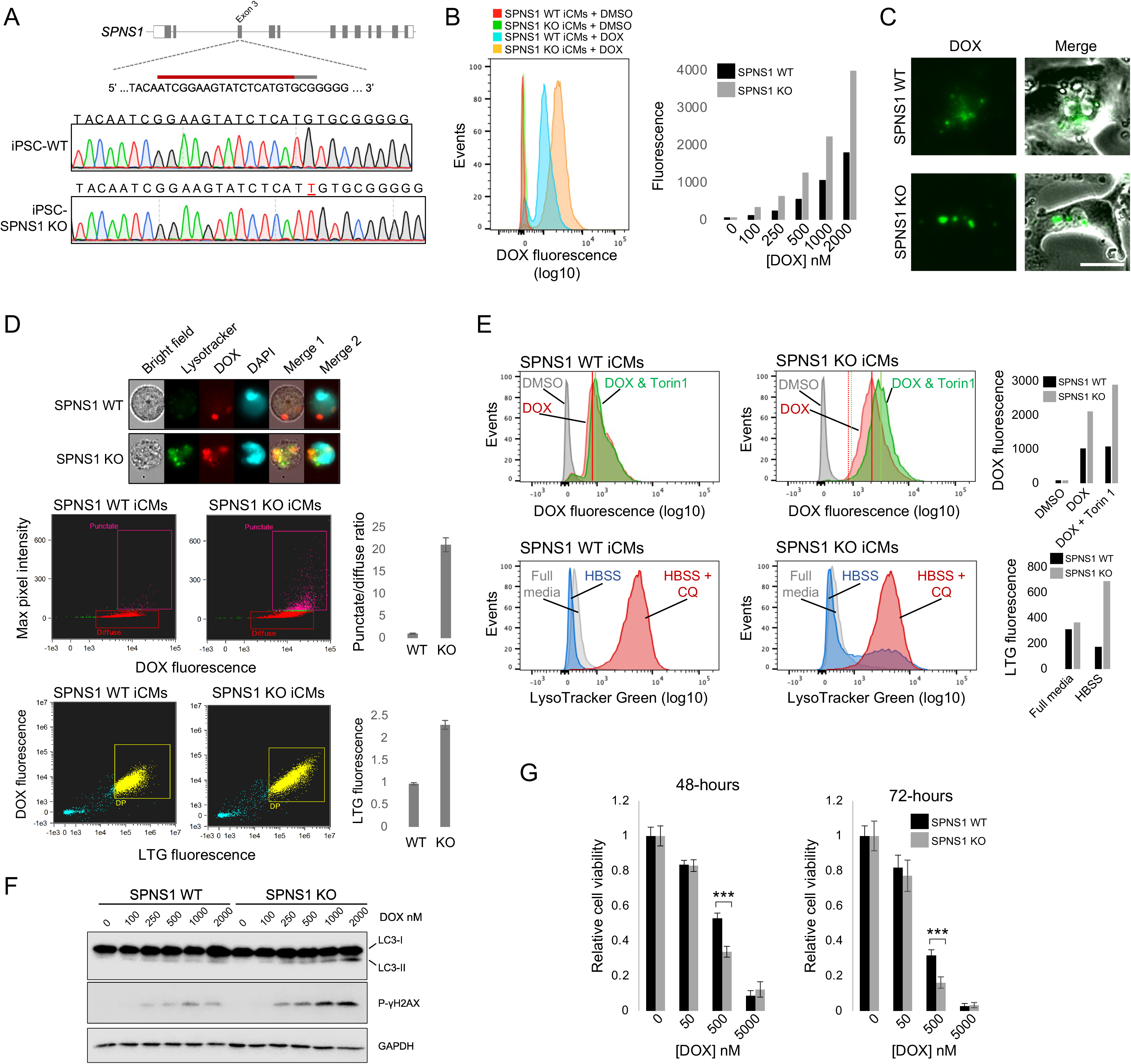
SPNS1 deficiency exacerbates DOX accumulation and toxicity in iPSC-derived cardiomyocytes. (A) Schema of SPNS1 gene structure, wildtype genomic DNA (gDNA) sequence, region targeted by sgRNA, and Sanger traces of SPNS1 wildtype (WT) and SPNS1 knockout (KO) iPSCs. (B) Flow cytometry analysis of DOX accumulation in SPNS1 WT and SPNS1 KO iPSC-derived cardiomyocytes (iCMs). Representative flow cytometry histogram (left) and background-corrected median fluorescence intensities following incubation with specified concentrations of DOX (right). (C) Immunofluorescence microscopy of DOX accumulation in SPNS1 WT and SPNS1 KO iCMs. Scale bar: 40μm. (D) Image cytometry analysis of DOX localization in SPNS1 WT and SPNS1 KO iCMs. Top panel shows representative image captures. Middle panels show DOX fluorescence versus max pixel intensity. Max pixel intensity is proportional to the degree of aggregation, visible as puncta. Bar charts shows relative amounts of DOX puncta in SPNS1 WT and SPNS1 KO iCMs. Bottom panels show LysoTracker Green fluorescence versus DOX fluorescence. (E) Top panels display effect of Torin 1 supplementation (50nM) on DOX accumulation in SPNS1 WT and SPNS1 KO iCMs. Bottom panels show effect of Hank’s balanced salt solution (HBSS) and chloroquine (CQ, 25μM, as a control for lysosome accumulation) on lysosome abundance in SPNS1 WT and SPNS1 KO iCMs. Bar charts show median fluorescence intensity from accompanying histograms. Data is from a single experiment but three assays using separate batches of iCMs yielded equivalent results. (F) Protein levels of LC3 and phosphorylated H2AX in SPNS1 WT and SPNS1 KO iCMs following treatment with specified concentrations of DOX for 24-hours. (G) Cell viability in SPNS1 WT and SPNS1 KO iCMs treated with specified concentrations of DOX for 48- and 72-hours. Cell viability was measured via alamarBlue assay (see Methods). \*\*\**P*<0.0001 (unpaired 2-tailed t test)

## Discussion

Here we use CRISPR/Cas9 screens to spotlight genes, proteins and pathways implicated in two events central to doxorubicin (DOX) cardiotoxicity. We first used survival-based screens to identify factors involved in DOX-mediated cell death followed by accumulation screens to discover factors that regulate DOX uptake and clearance. Over 270 drug-gene interactions were identified, underscoring the regulated nature of these important events. Follow-up studies revealed that retinoic acid receptor-*α* (RARA) status is a targetable determinant of DOX-mediated cell death and that Sphingolipid Transporter 1 (SPNS1) and autophagic lysosome reformation facilitates cellular clearance of DOX.

Anthracyclines comprise a group of highly effective chemotherapeutic reagents used for the treatment of various hematologic and solid tumor malignancies, including breast cancer, lymphoma, leukemia, and sarcoma. Despite critical survival gains, anthracyclines such as DOX elicit severe cardiotoxic side effects. The underlying pathological mechanisms are largely undefined and therapeutic options extremely limited. DOX adversely affects myriad crucial cellular processes (41). Given extensive cross-talk between these processes, parsing causal (i.e., driving) mechanisms from coincidental or associated events remains challenging. The current study attempted to do so by functionally interrogating genetic factors involved in DOX-mediated cell death and DOX accumulation.

In our cell death screens, canonical apoptosis factors were, as anticipated, amongst the strongest hit genes. Disruption of Apaf1 and Casp9, which form the apoptosome (42), desensitized immortalized cardiomyocytes, iPSC-derived cardiomyocytes (iCMs), and breast cancer cells to DOX toxicity, underscoring the role of regulated cell death in DOX cytotoxicity. Our screens also highlighted the role of multi-drug resistance proteins in buffering against DOX toxicity. ABCB1B (HL-1 screen) and ABCC1 (AC16 and iCM screens) loss-of-function correlated with increased DOX toxicity. Exactly how ABCC1 and ABCB1B facilitate DOX clearance is not known and requires further examination. The most significantly depleted sgRNAs in the iCM screen were in the RARA gene. This tallies with independent studies linking RARA status to cardiac health. For example, heart failure is associated with diminished RARA signaling (43) and vitamin A deficiency worsens maladaptive cardiac remodeling following myocardial infarction (44). Additionally, in a mouse model of myocardial infarction, cardiomyocyte-specific ablation of retinoic acid signaling increases infarct size and apoptosis (45). In our study, RARA activation using Tamibarotene (TBT) reduced DOX toxicity, although the protective mechanism is unclear. In RNA-Seq studies, a striking increase in mitochondrial gene expression in iCMs treated with TBT plus DOX compared with iCMs treated with DOX alone was observed. TBT alone did not induce such changes meaning it may buffer against DOX-mediated repression of mitochondrial genes. Whether this represents a causal mechanism or a by-product of improved cell health requires investigation. In support of the latter, the RARA target gene, Cytochrome P450 Family 26 Subfamily B Member 1 (CYP26B1), was the most significantly up-regulated transcript by TBT (log_2_FC=8.1, *P*=1.5×10^-21^, FDR<0.001) and was also markedly up-regulated by DOX alone (log_2_FC=6.3, *P*=5.6×10^-29^, FDR<0.001). It is tempting to speculate that CYP26B1 promotes DOX metabolic clearance and that pre-induction by TBT primes cardiomyocytes, expediting clearance and alleviating toxicity. A recent study by the Burridge group (46), informed by a prior GWAS of pediatric oncology patients (47), demonstrated that a common non-synonymous coding variant in the retinoic acid receptor-γ (RARG) gene (S427L) sensitizes human iPSC-derived cardiomyocytes to DOX toxicity, and activation of RARG using a small molecule (CD1530) conferred protection against DOX cardiotoxicity *in vivo*. It is possible that the protective effect of TBT results from preferential activation of RARG rather than RARA. Although TBT is reported to bind with stronger affinity to RARA than RARG, specificity will need to be empirically tested using RARA and RARG single and double knockout iCMs. It will also be important to test if simultaneous activation of RARA and RARG, via TBT and CD1530 respectively, imparts greater protection against DOX than either agonist alone. RARA loss-of-function had an opposing effect in our exploratory screen and confirmatory screen. This may be due to different genetic backgrounds, epistatic effects, or other factors. Notably, the allelic effect of the RARG rs2229774 polymorphism was also discordant in GWAS of anthracycline cardiotoxicity in breast cancer patients (48).

DOX has a propensity to accumulate within cardiomyocytes but mechanisms that drive uptake as well as promote clearance are poorly understood. We exploited the fluorescent properties of DOX to screen, via flow cytometry-activated cell sorting (FACS), for genes involved in its accumulation within cardiomyocytes. Interestingly, the quantity of hits and effect sizes in the low DOX uptake bin were relatively small. An explanation, consistent with previous studies (49), is that carrier-mediated DOX transport is negligible and that uptake occurs mainly via passive lipoidal diffusion. This model contrasts that proposed by Huang and colleagues (50), which supports active transport (via OCT3, which our screen did not discover). Further examination of DOX uptake routes is warranted. Irrespective of entry route, the hydrophobic nature of DOX enables passive diffusion into the lysosome compartment. Upon entry, the acidic lumen protonates DOX, preventing return to the cytosol, a process referred to as cation trapping. What is the fate of DOX following lysosomal sequestration? We observed DOX fluorescence in iCMs for several weeks after acute exposure, suggesting cardiomyocytes have limited capacity to clear DOX. This may explain why cumulative dose is an important risk factor for DOX cardiotoxicity. However, the fact that ABCC1 loss-of-function causes increased DOX accumulation implicates an efflux system and investigating how this is accomplished will be necessary next step.

A limitation of our study relates to physiological relevance of the screening system. Although we validated the effect of RARA (cell death screen) and SPNS1 (DOX accumulation screen) using iCMs, genome-wide screening in primary cardiomyocyte cultures, iCMs, and/or whole heart *in vivo* will be important if genetic modifiers are to be probed *en masse* in a physiologically relevant manner. A genome-wide screen of DOX cytotoxicity in iCMs was recently described (51) but statistics (specifically, adjusted *P* values) were unavailable so a formal head-to-head comparison was not performed.

Survival-based CRISPR screens rely on selective pressure to promote proliferation of some cells and slowing it in others. Given the low rate of mitosis in iCMs, capacity to identify enriched sgRNAs is weakened and a greater reliance is placed on depletion. Hence, it will be important to employ growth-independent methods, such as CROP-Seq (52), which could leverage the striking change in gene expression following DOX treatment as a screening readout. Knowles and colleagues recently showed DOX sensitivity stems from the cardiomyocyte’s in/ability to mount a robust transcriptional response, which is determined by genetic background (53). Hence, CRISPR screens across different genetic backgrounds (e.g., iCMs from African Americans versus European Americans) and disease contexts (e.g., iCMs derived from patients who received DOX and developed cardiotoxicity versus those who received DOX without ill-effect) to discover core and context-dependent liability genes (54) will also be informative.

A caveat of fluorescence-based screening is that perturbations which significantly modify cellular pH or lead to accumulation of auto-fluorescent molecules such as heme can cause false positives. UROD was joint-highest hit in our accumulation screen but was a false positive (data not shown), as was also the case in a fluorescence-based autophagy screen by Shoemaker and colleagues (55).

We established via quantitative and qualitative assays that SPNS1 loss-of-function exacerbates DOX accumulation and toxicity but the underlying mechanism requires further examination. A possible model is that elevated lysosome content in SPNS1 KO cells creates an expanded acidic compartment into which DOX can diffuse, promoting hyperaccumulation. This blocks autophagic flux, generating ROS and driving cardiotoxicity. Independently of these events, it is feasible that lysosomal accumulation of DOX elicits a stress response analogous to cholesterol (56), which may be therapeutically targetable. Indeed, we observed increased DNA damage in SPNS1 KO cells. Therefore, transient stimulation of ALR may be beneficial upon DOX administration.

The binary nature of CRISPR KO screening does not inform how subtle changes gene in expression alter phenotypes. Interestingly, alterations in SPNS1 transcript abundance are associated with body mass index (57), a trait which correlates with anthracycline cardiotoxicity risk (58). Hence, it will also be necessary to determine if physiologically-relevant changes in SPNS1 expression affect DOX accumulation and cytotoxicity.

## Methods

### CRISPR/Cas9 knockout screens

To establish an HL-1 cell line expressing Cas9, lentiCas9-Blast (Addgene plasmid #52962) was transfected into Lenti-X 293T cells (Takara) with pMD2.G (Addgene plasmid #12259) and psPAX (Addgene plasmid #12260). Viral-containing supernatant was added to HL-1 cells and Cas9-expressing cells were isolated by culture in blasticidin (10μg/ml)-containing media for 2 weeks.

For HL-1 cell death screens, mouse GeCKOv2 CRISPR knockout pooled library (Addgene #1000000053,18), pMD2.G and psPAX2 were transfected into Lenti-X 293T cells. Three days post-transfection, supernatants were pooled, incubated with Lenti-X solution (Takara) and concentrated by centrifugation. Viral titer was determined by transducing HL-1 cells with serial dilutions of lentivirus and measuring puromycin resistance. For AC16 cell death screens (and DOX accumulation screens, see below), the same procedure was conducted to create and quantify the one-vector format Brunello human CRISPR knockout pooled library (Addgene #73179,59).

HL-1-Cas9 cells and AC16 cells were transduced with GeCKOv2 and Brunello libraries, respectively at ~300x coverage (i.e., each sgRNA was delivered into ~300 cells) and a multiplicity of infection (MOI) of ~0.2 to ensure most transduction events involved a single lentivirus particle. Two days post-transduction, cultures were switched to puromycin (1μg/ml for HL-1 cells, 2μg/ml for AC16 cells)-containing media and maintained therein for 1 week to select for stably-transduced cells.

For HL-1 and AC16 cell death screens, mutagenized cells were seeded across multiple 10cm plates (~10M cells/plate) and randomly assigned to either an experimental or a control group. Experimental cultures received media containing DOX (Cell Signaling) at 1μM whilst control cultures received media containing DMSO. Library coverage of ~300x was maintained throughout the experiment. After 48-hours, remaining cells were harvested, counted, and frozen at −80°C until needed.

For DOX accumulation screens, mutagenized AC16 cells were incubated in standard growth media containing 1μM DOX for 24-hours. Cells were detached via trypsin and sorted into high DOX content (top 15%) and low DOX content (bottom 15%) sub-populations using a BD FACSAria II. A population of mutagenized cells (~30M) which were treated with DOX but not sorted was also prepared. Library coverage of ~300x was maintained at all steps. Cells were frozen and stored at −80°C until needed.

### sgRNA library preparation and data analysis

Genomic DNA was isolated using Quick-DNA Midiprep Plus Kit (Zymo). sgRNA cassettes were amplified using round 1 (R1_F and R1_R in Table 1) primers. Stagger sequences were added to round 1 amplicons using round 2 (R2_F1-9 and R2_R in Table 2) primers, and barcodes were added to round 2 amplicons using round 3 (R3_F and R3_R in Table 2) primers. Equimolar quantities of each amplicon were pooled and sequenced on an Illumina NovaSeq. Resulting FASTQ files were quality-trimmed and sample comparisons were conducted using MAGeCK as previously described (19).

**Table 1.**
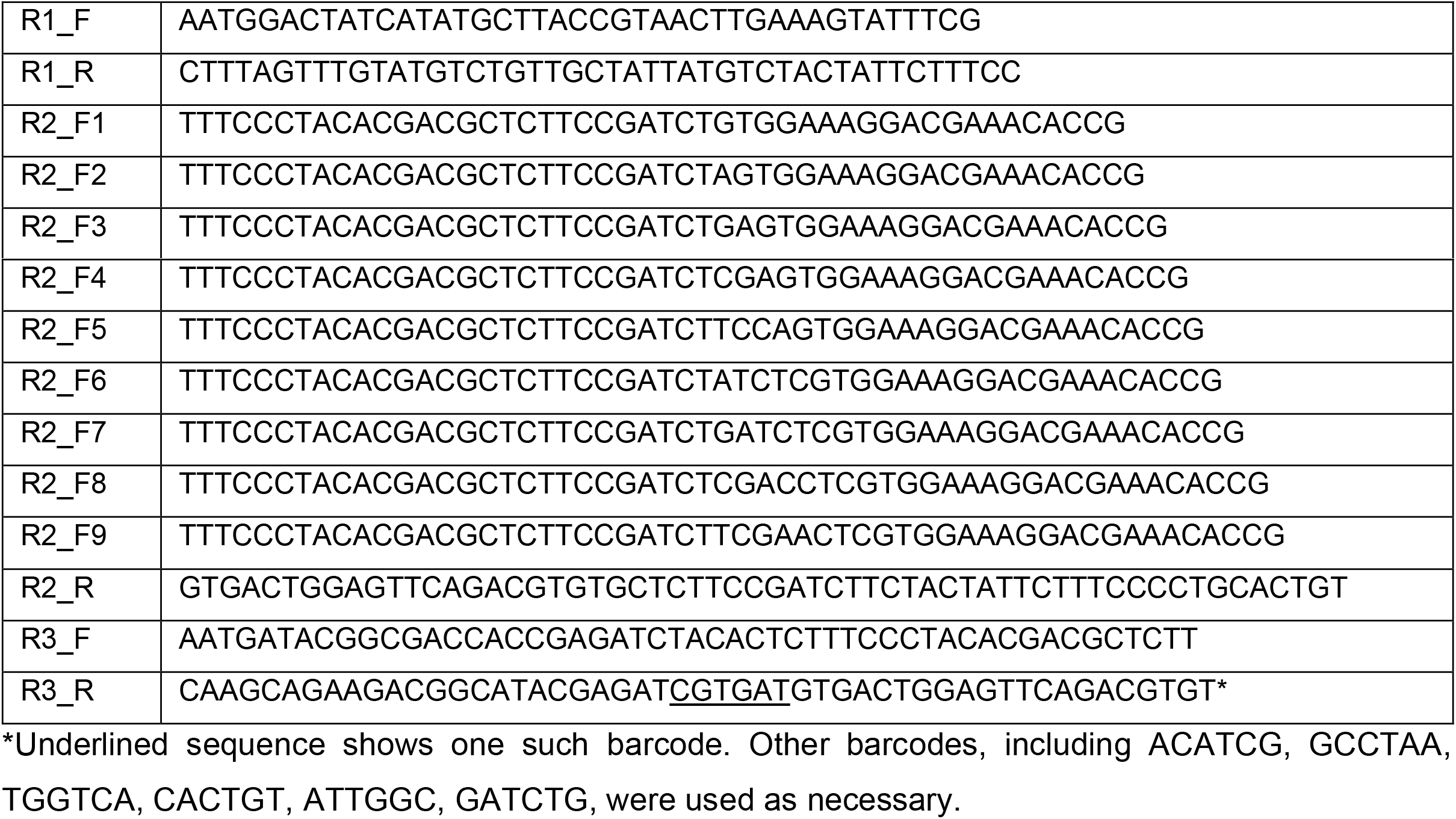
HL-1 screen primers.

**Table 2.**
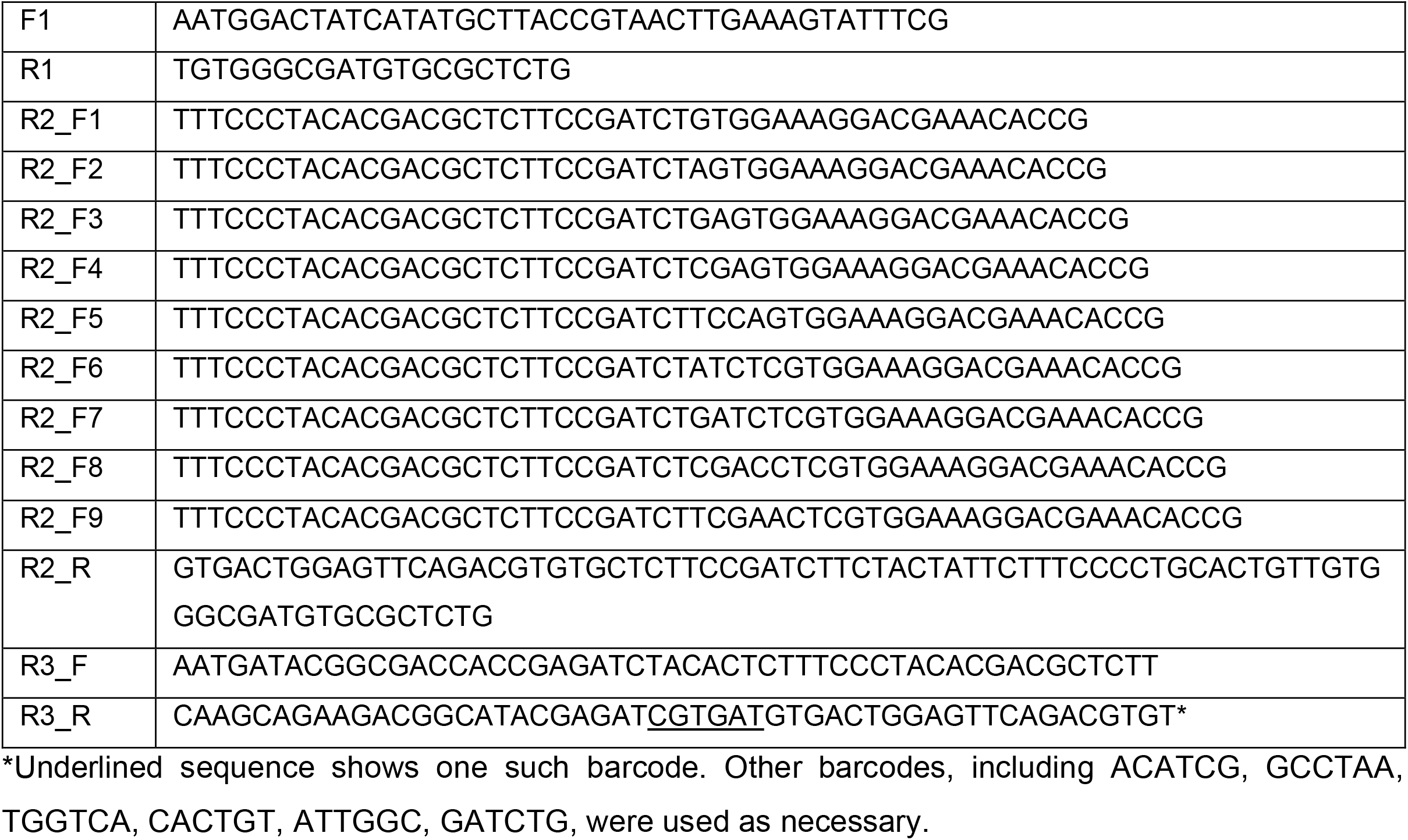
AC16, iPSC-CM, and MCF-7 screen primers.

### Generating knockout cell lines and data validation

Multiple genes from the HL-1 whole-genome CRISPR/Cas9 knockout (KO) screen were selected for validation. Target sites were identified using CHOPCHOP (60) and prioritized based on 5’ positioning, predicted cleavage efficiency, and specificity. Oligonucleotides were hybridized and cloned into lentiCRISPR v2 (Addgene plasmid # 52961). Viral supernatants, prepared as described above, were applied to HL-1 cells. Following 7-10 days in media containing puromycin, genomic DNA was isolated and indel frequency was estimated using TIDE (61). Cell lines harboring >60% disruptive mutations were retained.

For DOX toxicity assays, cells were seeded in 96-well plates. Each plate consisted of wild type (i.e., untransduced) cells, cells transduced with a non-targeting sgRNA, and KO cells. The following day, cells received fresh media containing doxorubicin (0,0.05,0.5,5μM). After 3-days, cells were processed with alamarBlue (Thermo) and fluorescence, which reflects metabolic flux and hence viability, was measured using a BioTek Synergy microplate reader. Data were analyzed in Microsoft Excel.

### Targeted CRISPR/Cas9 KO library design and synthesis

A custom CRISPR/Cas9 KO library targeting the highest scoring hits from the HL-1 screen and the AC16 screen was generated. The targeted library comprised 7820 sgRNAs targeting 952 genes (8 sgRNAs per gene) and 200 non-targeting sgRNAs. Oligonucleotides were synthesized (Agilent) and Gibson-cloned into lentiCRISPR v2 and analyzed as described previously (62,63). Briefly, library coverage (99%) and sgRNA skew factor (4.3) were well within recommended limits.

### Maintenance of iPSCs and cardiomyocyte differentiation

iPSCs were maintained in a feeder-free format on Geltrex-coated 6-well plates using StemMACS iPS-Brew XF media (Miltenyi Biotec) with daily feeding. Cells were passaged (1:12) every 3-4 days using StemMACS Passaging Solution XF (Miltenyi Biotec) and Y-27632 dihydrochloride (Santa Cruz) to reduce cell death. Pluripotency was assessed every 8 weeks via immunofluorescence detection of NANOG (Cell Signaling, 4903) and SSEA4 (Cell Signaling, 4755). iPSCs were differentiated into cardiomyocytes (iPSC-CMs) via the Wnt/β-catenin modulation protocol (25), with a matrix overlay to enhance epithelial-to-mesenchymal transition (64), and lactate selection to promote maturation (26). Purity was assessed by measuring TNNT2 expression by flow cytometry and was consistently >90%.

### Targeted CRISPR/Cas9 KO screens

A healthy donor-derived iPSC line (SV20,24) was transduced with lentivirus at ~1000x coverage and an MOI of ~0.3 with the targeted CRISPR/Cas9 KO library. Following selection in puromycin media (1μg/ml pulsed for 3 days), cells were expanded, assessed for pluripotency via immunofluorescence detection of NANOG (Cell Signaling, 4903) and SSEA4 (Cell Signaling, 4755), and cryopreserved. Following differentiation, mutant iPSC-derived cardiomyocytes (iCMs) were seeded across multiple 10cm plates and randomly assigned to an experimental or control group. The experimental group received fresh media containing 250nM DOX for 5 days whilst the control group received fresh media containing DMSO only. Genomic DNA was isolated and sequencing libraries were prepared and analyzed as described above. Each gene was targeted by 8 sgRNAs. In many cases, we observed significant heterogeneity in sgRNA abundance for a given gene. This was observed in both biological replicates in the iPSC and iPSC-CM screen indicating variable sgRNA activity was the cause. For this reason, we applied a modified inclusion threshold to select modifier genes. A gene was defined as a hit when ≥3 of the sgRNAs targeting the gene exhibited a ≥2-fold change in abundance in the DOX treatment group compared with the DMSO-treatment group with the sgRNA showing a concordant change (i.e., either enriched or depleted but not both). Using this strategy, we nominated 46 high confidence gene-drug interactions.

### RARA experiments

For RARA knockout, CRISPR target sites in the RARA gene were identified using CHOPCHOP (60). CRISPRs were designed against two adjacent target sites and cloned into pX458 (65). Cloning insertion was confirmed by Sanger sequencing and endotoxin-free plasmid preparations were generated. Plasmid DNA was transfected into hB53 iPSCs via nucleofection. Cells were sorted based on GFP expression the following day and seeded at low density (1000-3000 cells) in 10cm plates. Single colonies were picked, propagated, and genotyped by PCR and Sanger sequencing.

For RARA loss-of-function experiments, wild type and RARA KO hB53 iPSCs were differentiated into iCMs as described above. iCMs were seeded into 96-well plates (100K cells/well). After 72-hours, cells received media containing DOX (0.001,0.01,0.05,0.075,0.1,0.5,1,10μM). After 3-days, cells were processed with alamarBlue (Thermo) and fluorescence was measured using a BioTek Synergy microplate reader. Data were analyzed in Microsoft Excel.

For RARA gain-of-function experiments, wild type hB53 iCMs were seeded into 96-well plates and treated with media containing 1μM Tamibarotene (TBT) or DMSO. Following 24-hours, cells then received fresh media containing 1μM TBT or DMSO as well and DOX (0.001,0.01,0.05,0.075,0.1,0.5,1,10μM). After 3-days, cell viability was analyzed as above.

### RNA-Seq studies

Poly-A mRNA was prepared from day-30 iPSC-CM cultures treated with DMSO, DOX, TBT plus DOX, and TBT (4 biological replicates per condition) with RNeasy columns (Qiagen), followed by oligo-dT selection. Sample quality was assessed using the Agilent Bioanalyzer 2100. Library preparation was conducted with TruSeq RNA Library Prep Kit v2 chemistry, and sequencing was carried out on an Illumina HiSeq 2500. FASTQ reads were aligned to human genome (hg19) with STAR 2.5.2a (66). Transcript counts per million files were normalized by the Cox-Reid method and compared via PCA (FactoMineR) and quasi-likelihood F-tests in edgeR (67). Gene set enrichment analysis was conducted using clusterProfiler (68).

### SPNS1 knockout

An independent SPNS1 sgRNA was designed using CHOPCHOP (60). Forward and reverse oligonucleotides (CACCGAATCGGAAGTATCTCATGTG and AAACCACATGAGATACTTCCGATTC) were hybridized and cloned into lentiCRISPR v2 and lentivirus particles were generated as described above. Crude SNPS1 lentivirus preparation was transduced into AC16 cells and replaced with standard media 6 hours later. To control for possible confounding effects of lentivirus transduction, AC16 cells were transduced with lentivirus encoding a non-targeting sgRNA. Cells were maintained in puromycin-containing media for ~7 days, after which gene disruption was confirmed via Sanger sequencing and CRISPR Tide (61).

For SPNS1 knockout in iPSCs, endotoxin-free plasmid preparation was generated and used to create high titer lentivirus, which was then transduced into hB53 iPSCs. Cells were cultured in media containing puromycin (1μg/ml) for 1-week, after which single colonies were picked, propagated, and genotyped by PCR and Sanger sequencing.

### Flow cytometry and image cytometry analysis of DOX uptake

For conventional flow cytometry, cells were treated with DOX and other chemicals as described in the main text. Cells were washed with PBS, detached with trypsin, centrifuged, and washed again with PBS. After being centrifuged for a second round, cells were resuspended in ~500ul PBS and analyzed using a BD FACSCanto flow cytometer. Forward and side scatter plots were used to gate for single cells (20,000 cells per sample). Median fluorescence intensity at 585nm was measured following excitation with the blue laser (20,000 singlets per sample). FCS files were processed and analyzed using FlowJo (BD).

For image cytometry, cells were treated with DOX for 24-hours and then dispersed as described above. Vybrant DyeCycle Violet (ThermoFisher) and 75nM LysoTracker Green were added to the cell suspension, which was then incubated at 37°C for 30-minutes. Cells were analyzed on an Amnis ImageStream.

### Fluorescence microscopy

Timelapse fluorescence imaging of DOX uptake was carried out using a Nikon AX R confocal microscope fitted with an atmospheric control chamber (5% CO2 and 37°C). Cells were seeded in glass bottom dishes (Mattek) and the following day, received media containing 1μM DOX. Images (brightfield and 488nm laser) were acquired with a 40x objective every 20 minutes for 10 hours.

## Acknowledgements

We thank Andrea Stout and Xinyu Zhao from the CDB Microscopy Core for assistance with cell imaging studies, Richard Schretzenmair and all members of the Penn Cytomics and Cell Sorting Resource Laboratory for help with cell sorting and image cytometry, and Quentin McAfee and Sunhye Jeong for useful discussion. This work was funded by grant R35-HL145203 from the U.S. National Institutes of Health and by the Winkelman Family Fund in Cardiovascular Innovation.

---

**Supplementary Figure 1:**
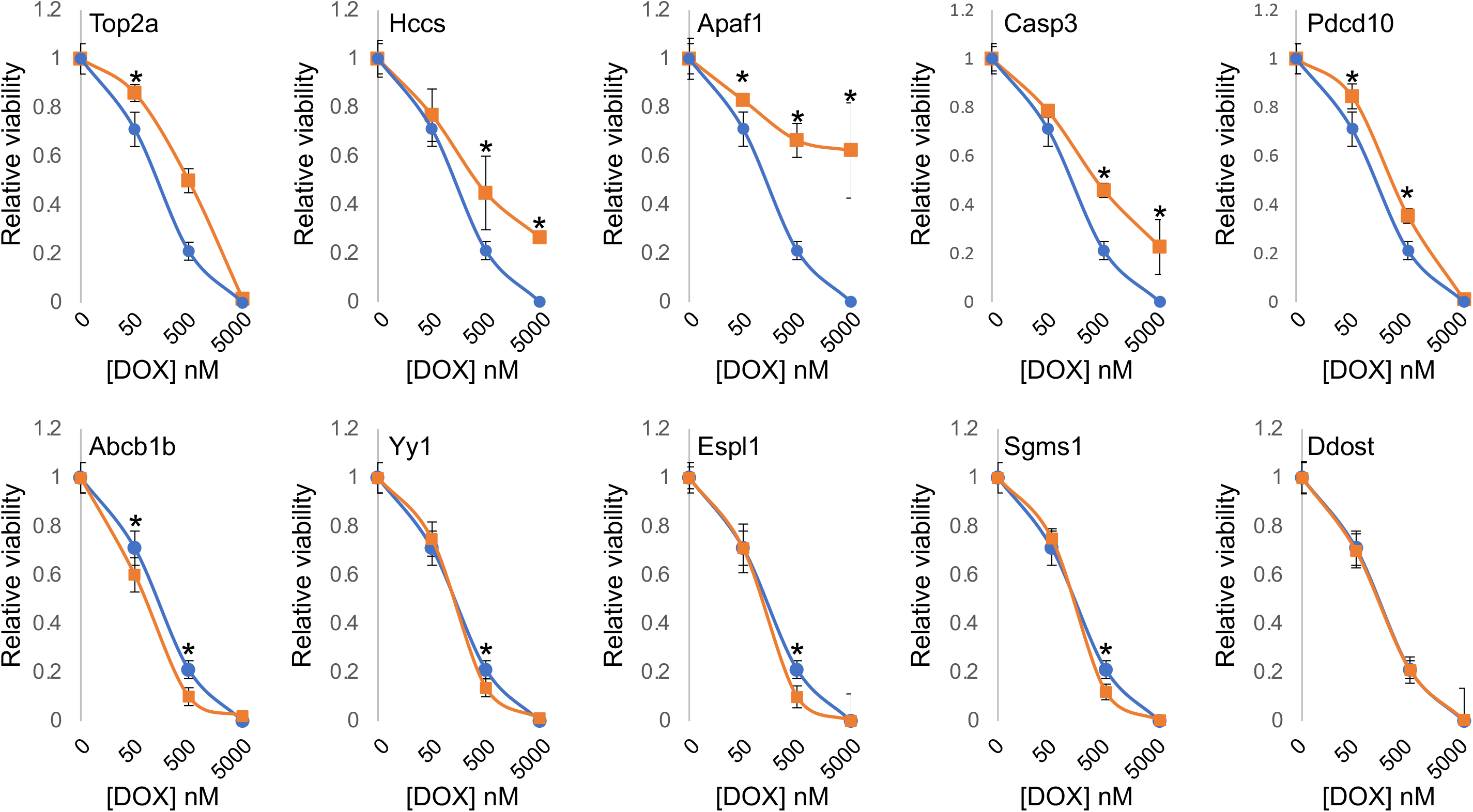
Validation of screening data. High scoring genes from HL-1 screen were targeted via CRISPR to generate independent bulk knockout (KO) cell lines. KO cells (orange data points/curve) and cells transduced with a non-targeting sgRNA (blue data points/curve) were treated with specified concentrations of DOX for 3 days and viability was measured via alamarBlue assay. Each data point is average and standard deviation of ≥6 biological replicates. *P<0.05, calculated by two-sided unpaired t test relative to HL1-con.

**Supplementary Figure 2:**
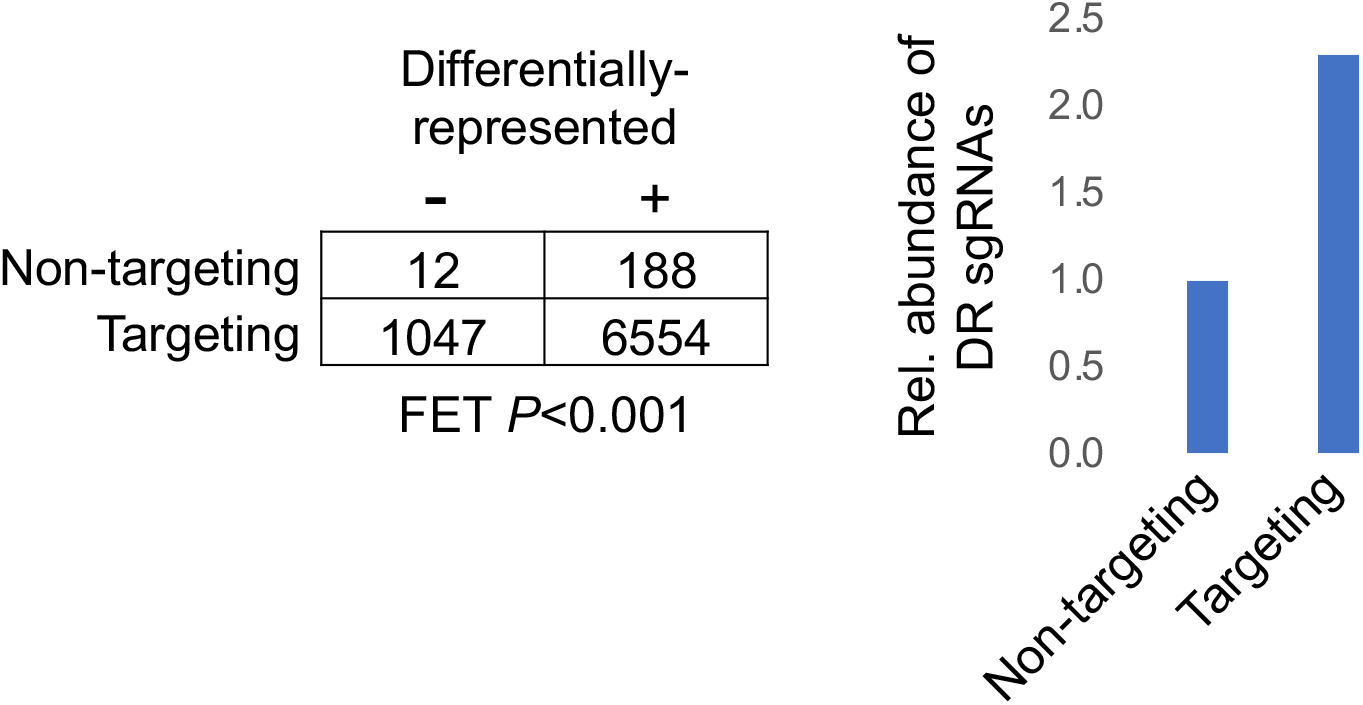
Contingency table-based comparison of targeting and non-targeted sgRNA representation in human iPSC-derived cardiomyocyte (iCMs) following DOX treatment. Left, contingency table and *P* value following Fisher’s exact test (FET). sgRNAs were classified as being differentially-represented (+) or not differentially-represented (-) if relative representation of the sgRNA was enriched or depleted by ≥2-fold in DOX-treated iCMs vs DMSO-treated iCMs. Right, relative abundance differentially-represented (DR) non-targeting and targeting sgRNAs. As expected, significantly more targeting sgRNAs were differentially-represented than non-targeting sgRNAs.

**Supplementary Figure 3:**
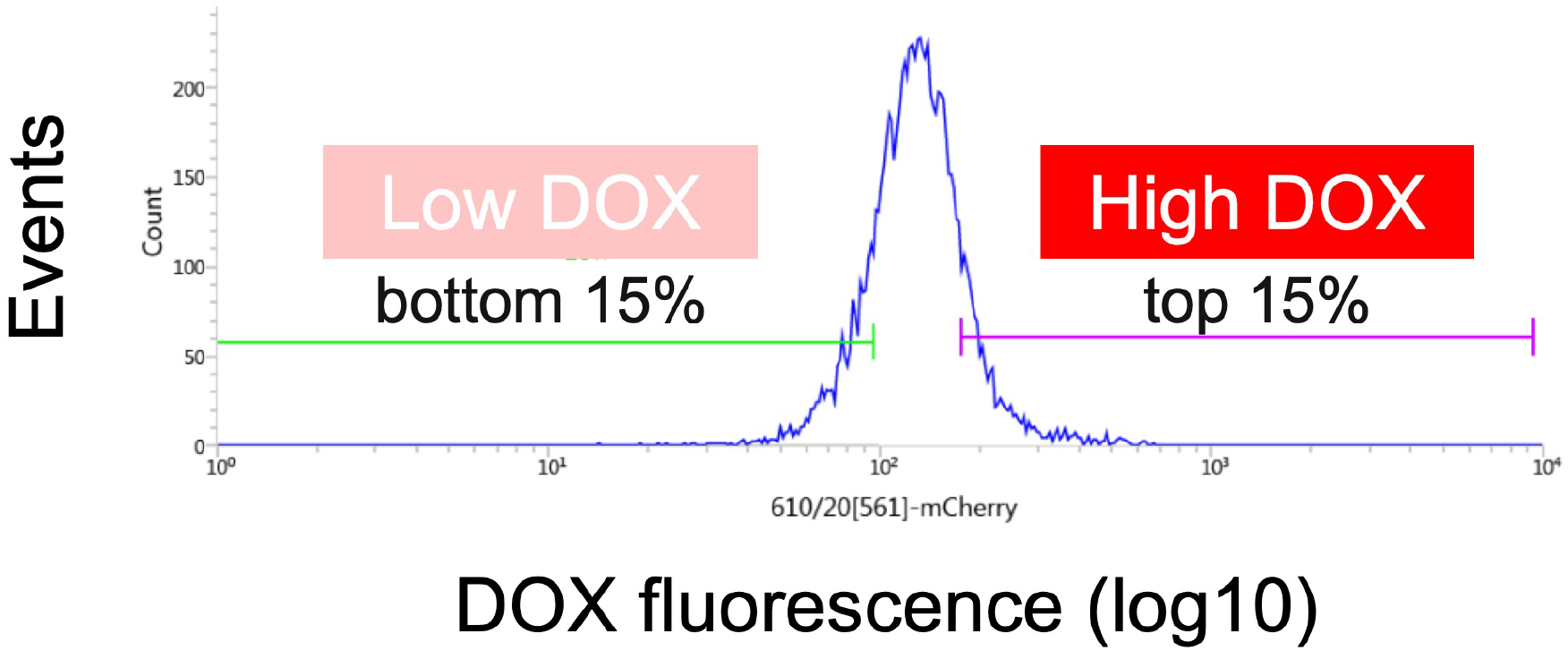
Histogram showing binning used for FACS-based isolation of low and high DOX content cells.

**Supplementary Figure 4:**
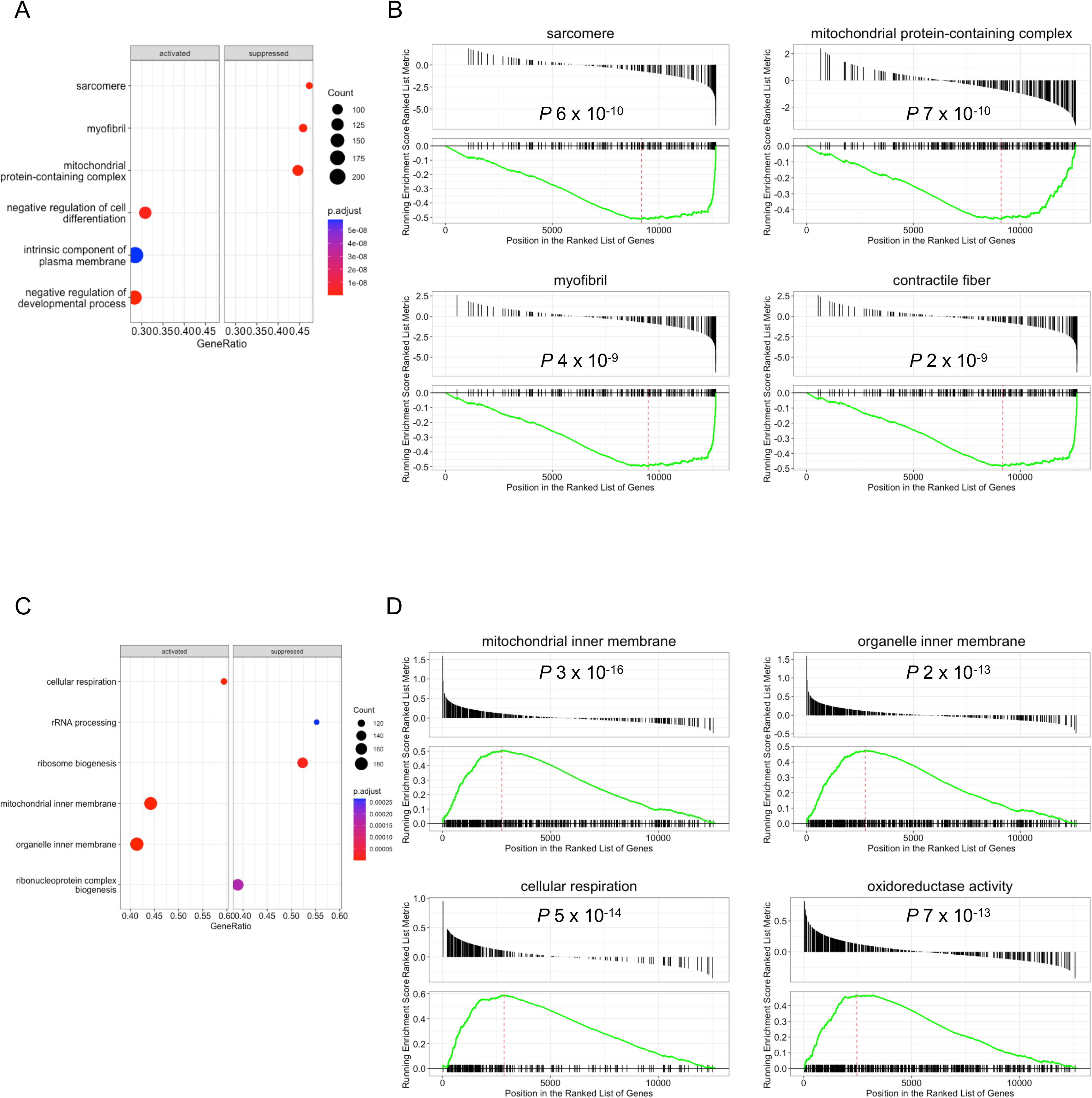
Gene set enrichment analysis of biological themes associated with doxorubicin (DOX) and Tamibarotene (TBT) treatment. Transcripts differentially-expressed (FDR<0.05) between the specified treatment groups were analyzed for association with gene ontology (GO) terms encompassing molecular functions, cellular components, and biological processes. (A) and (B) Analysis of GO terms differentially represented between DOX-treated iCMs and DMSO-treated iCMs. (C) and (D) Analysis of GO terms differentially represented between DOX & TBT-treated iCMs and DOX-treated iCMs. (A) and (C) Top terms in each comparison. (B) and (D) Enrichment profiles for the top four categories in each comparison.

**Supplementary Figure 5:**
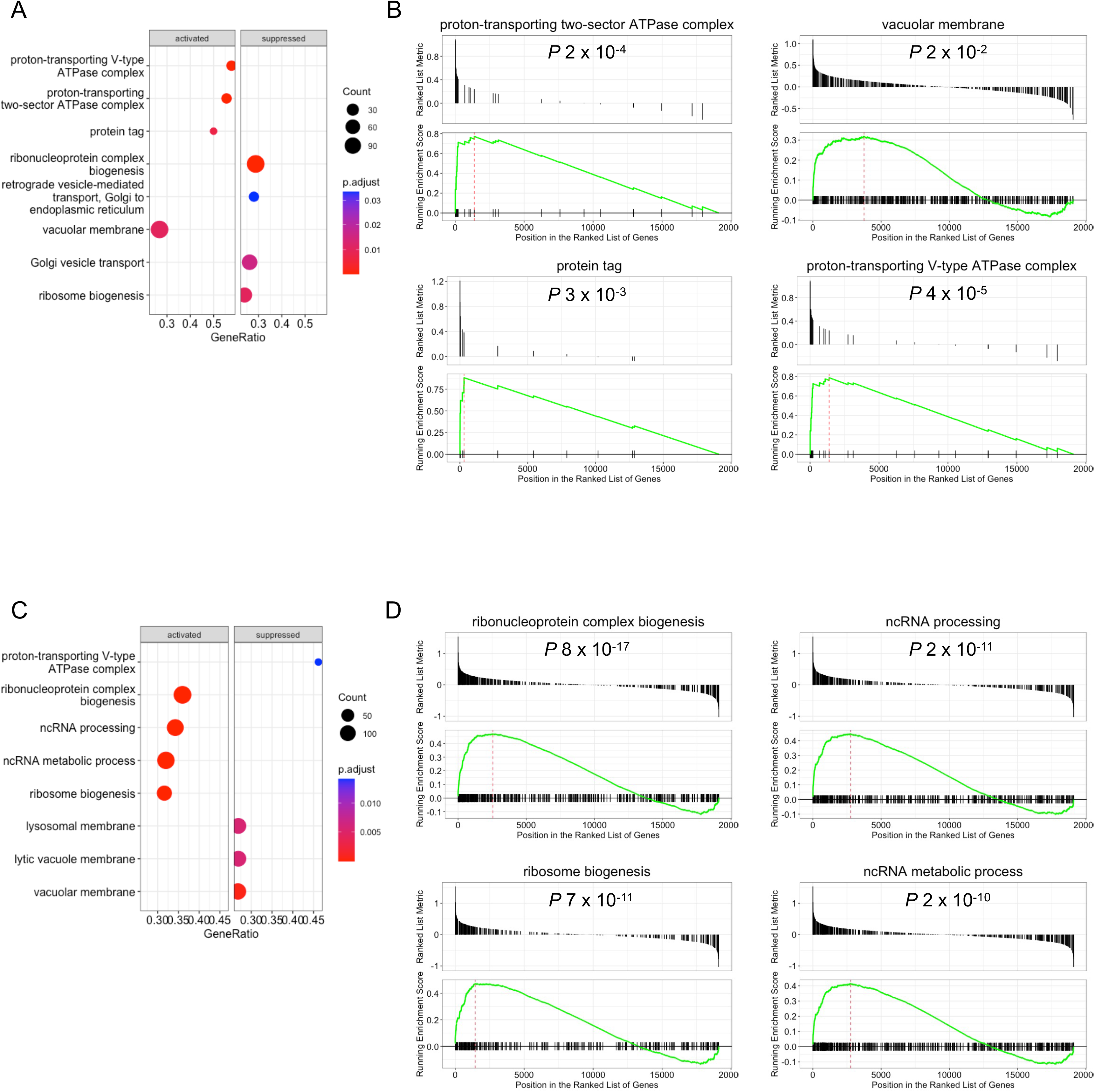
Gene set enrichment analysis of genes whose perturbation increases and decreases doxorubicin (DOX) accumulation in AC16 cells. (A) Top enriched and depleted (activated and suppressed, respectively) gene ontology (GO) terms in the high DOX bin. (B) Enrichment plots for the top four enriched GO terms in A. (C) Top enriched and depleted (activated and suppressed, respectively) GO terms in low DOX bin. (D) Enrichment plots for the top four enriched GO terms in C. Bonferroni corrected *P* values are shown for each GO term.

**Supplementary Figure 6:**
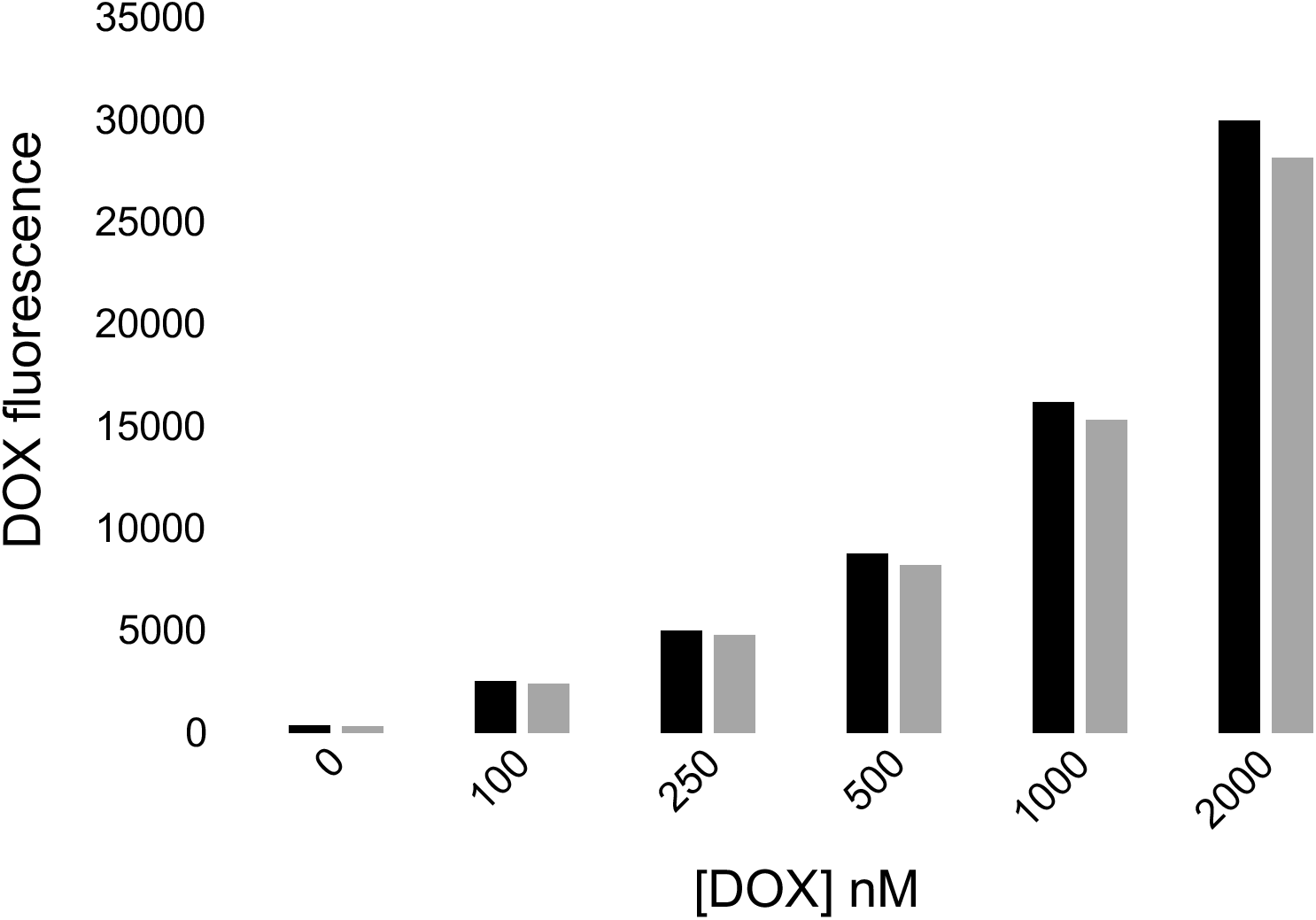
Doxorubicin accumulation in AC16 cells over-expressing Beclin-F121A and wild type AC16 cells. Black bars show doxorubicin (DOX) accumulation (1μM in standard growth media for 24-hours) in wild type cells, grey bars show accumulation in Beclin-F121A cells. Data are representative of three different experiments.

**Supplementary Figure 7:**
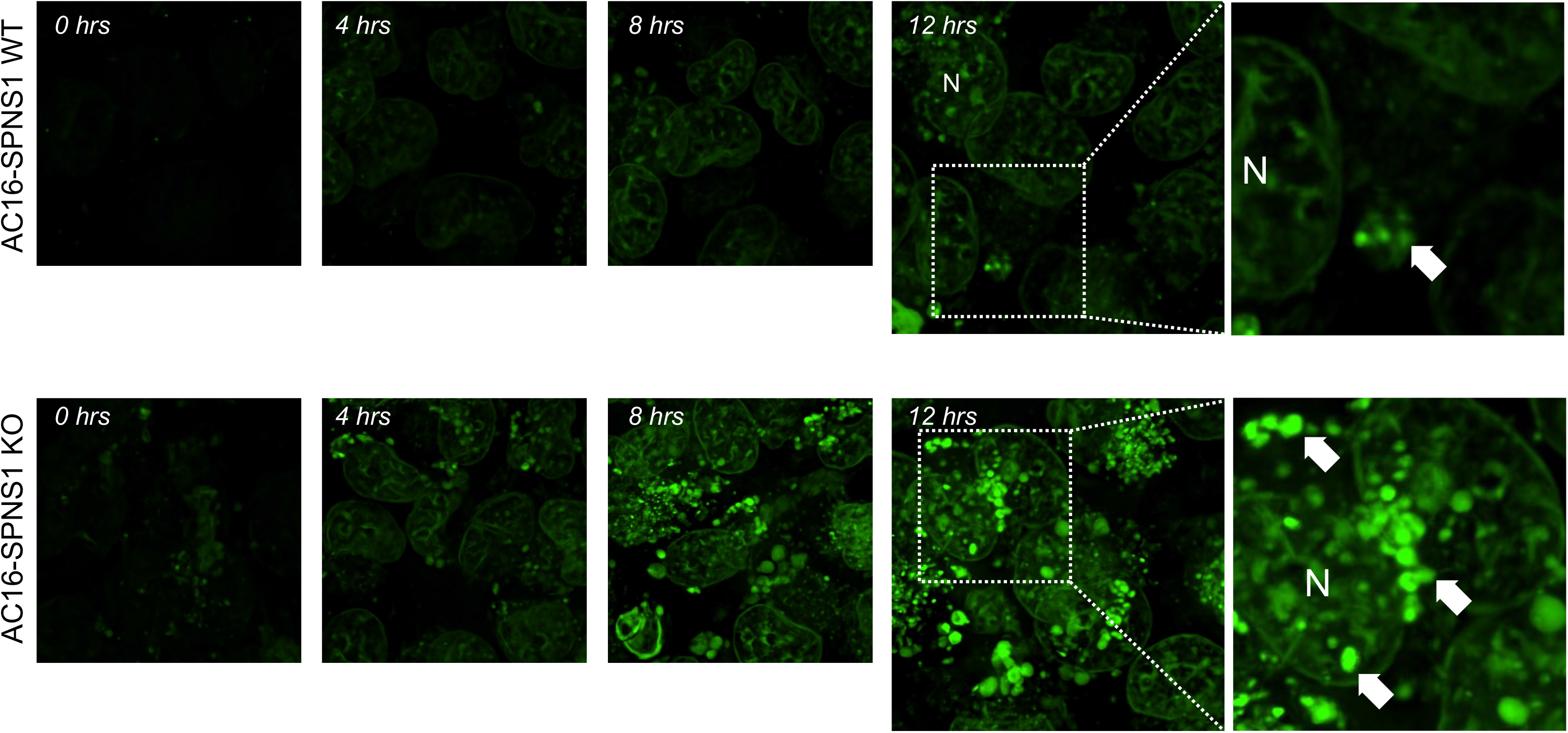
Time-lapse confocal microscopy of doxorubicin accumulation in SPNS1 wild type and SPNS1 knockout AC16 cells. Doxorubicin (DOX) was applied to wild type and SPNS1 knockout cells (1μM) in standard growth media and DOX uptake/accumulation was imaged every 20-min for a total duration of 12-hours. 0-,4-8,12-hour timepoints shown. DOX is pseudocolored green for clarity. Scale bar can be found in Figure 3. N, nucleus; arrows specify perinuclear DOX aggregates.

**Supplementary Figure 8:**
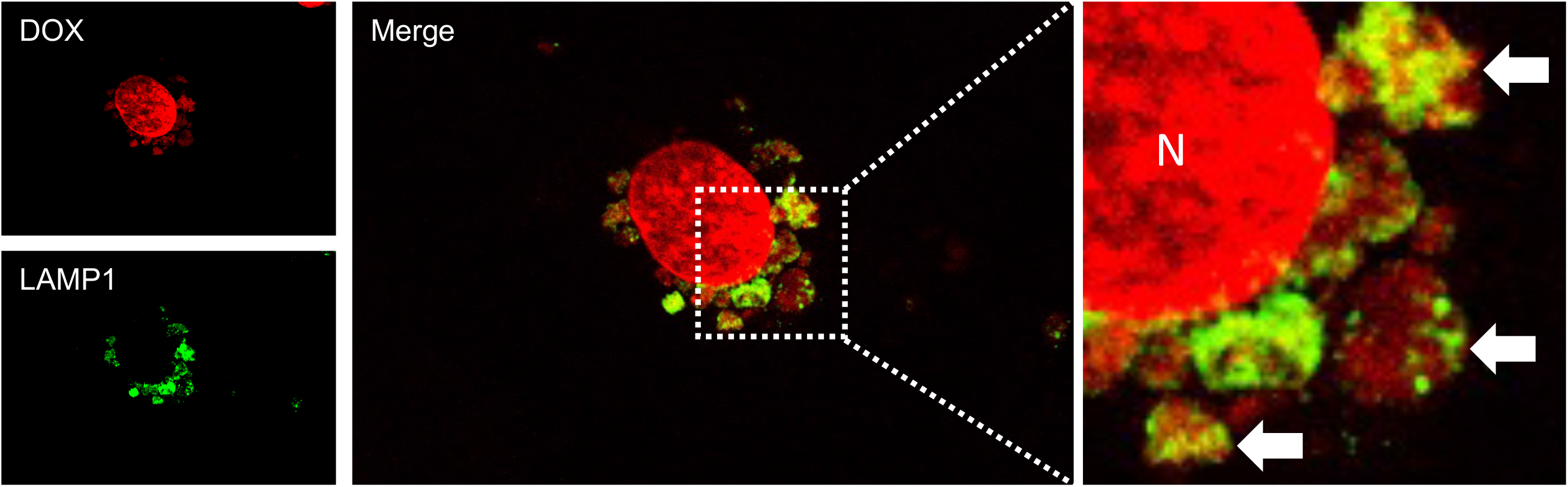
Lysosomal accumulation of doxorubicin in iPSC-derived cardiomyocytes. iPSC-derived cardiomyocytes (iCMs) were treated with 1μM doxorubicin (DOX) for 24-hours. DOX and LAMP1 (a lysosomal marker protein) distribution were investigated by confocal microscopy. Small (left-most) panels show DOX signal (pseudo-colored red) and LAMP1 (pseudo-colored green) in a representative iCM. The large (central) panel shows merge/co-localization between DOX and LAMP1. The right-most panel shows an enlarged, representative region of interest. N, nucleus. Arrows label perinuclear bodies positive for DOX and LAMP1.

## Notes

### Competing Interest Statement

The authors have declared no competing interest.

